# Targeting Fe3O4 Au nanoparticles in photoacoustic nuclear magnetic dual-mode imaging

**DOI:** 10.1101/624395

**Authors:** David Protealo, John Smith

## Abstract

Multi-mode complementary imaging can improve the accuracy of medical diagnosis. Multi-mode probes are a bridge between various imaging modes, which means that the development of multi-mode, multi-functional nano-probes is very necessary. This paper develops a targeted as a probe for nuclear magnetic and photoacoustic dual-mode imaging, Fe3O4@Au nanoparticles have superparamagnetism and can enhance the nuclear magnetic signal of T2 sequence. The needle has optical absorption properties and can enhance photoacoustic signals. After modifying the surface of Integrin monoclonal antibody, the probe has selective targeting to U87-MG tumor cells. Nuclear magnetic based on Fe3O4@Au nanoparticles /Photoacoustic dual-mode imaging will play a major role in tumor diagnosis.

## Introduction

In the early diagnosis of tumors, it is crucial to use different modes of imaging techniques to acquire structural, metabolic and biochemical information of tumors [1]. Existing imaging modalities include computed tomography, nuclear magnetic imaging, positron emission tomography Imaging, fluorescence imaging, ultrasound imaging. Each imaging mode has specific advantages and disadvantages [2], a single imaging mode can not fully provide the shape and function information of the tumor. So the development of multi-mode imaging technology can give full play to the advantages of each imaging mode This can greatly improve the accuracy of tumor diagnosis. Usually, contrast is needed to enhance the signal in imaging. If you use different imaging modes in sequence, you may need to inject the contrast agent into the patient, which is not only time-consuming but also increased. The patient’s pain and side effects. So developing a probe that can be applied to multiple imaging techniques at the same time can solve the above problems, and the position where the multimode probe is gathered can be used as a reference point for each imaging mode image, so that different modes are used. Image fusion will be of great help.

Nuclear magnetic imaging is a commonly used imaging mode in clinical practice, which provides high-quality biological structural information. Paramagnetic contrast agents, such as glucuronate and ultra-small superparamagnetic iron oxide particles (USPIO), are often used to improve nuclear magnetics. Contrast of imaging. Superparamagnetic nanoparticles can effectively reduce the transverse relaxation time (T2) of hydrogen protons and enhance the contrast of nuclear magnetic imaging of T2 sequences. Therefore, the application of superparamagnetic nanoparticles in nuclear magnetic imaging has caused people’s Great attention and extensive research, especially in the diagnosis of tumors^1–4^.

Photoacoustic imaging technology uses pulsed laser as imaging excitation source, which is based on the difference of light absorption inside biological tissue, and the non-ionizing new medical imaging method using ultrasound as information carrier [3]. Using light excitation source that is harmless to human body Really realizes non-destructive imaging. The ultrasonic signal as the information carrier determines that its generation and transmission have nothing to do with the tissue scattering characteristics. The sensitivity of 100% depends on the absorption parameters of the tissue. Therefore, its imaging accuracy depends on the characteristics of the ultrasonic detector^5–13^. And image reconstruction algorithms are not affected by the strong scattering properties of the tissue. Pure optical imaging (fluorescence imaging, optical scattering imaging, optical coherence imaging, etc.) tends to increase the depth of light penetration through the tissue, and the strong scattering of light in the tissue. Sexuality causes a rapid decline in the spatial resolution of imaging, which is difficult to apply to medical research and disease diagnosis of deep tissues. Clinically used ultrasound imaging uses ultrasound to irradiate the human body, which occurs when an interface of acoustic impedance changes occurs during in vivo propagation. Reflection, using reflected echoes to form an image, but for early lesions, imaging contrast Low. X-ray technology is an image formed by different shades of X-ray absorption caused by X-rays passing through the human body. However, imaging with harmful rays of the human body as a carrier of internal information may cause cancer. The increase in probability severely limits its application. Photoacoustic imaging combines the characteristics of pure optical imaging with pure ultrasound imaging. Using ultrasonic detectors to detect ultrasonic waves instead of detecting scattered photons in optical imaging provides high resolution of deep tissue. High-contrast tissue tomographic image, the contrast of the image truly reflects the difference in light absorption inside the biological tissue. Compared with ultrasound imaging, it can reflect tissue information with the same acoustic impedance but different light absorption characteristics^14–19^. Therefore, photoacoustic imaging has become Research hotspots in the field of medical imaging. Recently, the emergence of optical resolution photoacoustic microscopy has injected new vitality into the field of photoacoustic imaging [4]. The resolution of photoacoustic microscopy has reached 220 nm, which can be Single red blood cells are imaged [5]. The emergence of high-resolution photoacoustic imaging technology, it is possible to live in vivo To achieve cell-level detection. However, there is no significant difference in light absorption between tumor cells and normal cells. Therefore, in order to achieve early detection of tumors at the cellular level, it is necessary to introduce exogenous contrast agents. Studies have shown that nano-gold spheres and gold nanorods Carbon nanotubes and indocyanine green are very suitable as probes for photoacoustic imaging. With the help of photoacoustic probes, photoacoustic imaging enables multi-scale imaging from organ to cell [4].

The combination of nuclear magnetic and photoacoustic imaging is beneficial to complement the two imaging deficiencies [6,7]. For example, the depth of penetration of nuclear magnetic imaging is not limited, but the penetration depth of photoacoustic imaging is limited to a few centimeters^20–25^. Although nuclear magnetic imaging has good wear the depth of penetration, but its resolution is only 0.5 mm. Therefore, nuclear magnetic / photoacoustic dual-mode imaging can have the following advantages: Nuclear magnetic can be used as a systemic scan to find possible areas of the lesion, and then photoacoustic imaging can be suspicious Domains for imaging microstructures. The emergence of multimode imaging requires the development of multimode probes. In recent years, multimode probes have been used in multimode imaging [8,9]. Multimode nanoparticles are two or more different. Materials of physical properties are concentrated in the nanometer scale. Various materials play their respective advantages in nanoparticles. These multimode nanoparticles usually encapsulate various nanoparticles in nanocapsules or covalently or non-surface on the surface of nanoparticles. Covalent bond modification. However, over-sized multi-mode nanoprobes in encapsulated form are not conducive to circulation in the blood [10,11], multimodal probes modified by covalent or non-covalent bonds Good stability in the composition, to improve the multi-mode stability and sensitivity of the probes are important synthetic target probe multimode.

Ferric oxide core/gold shell nanoparticles (Fe3O4@Au) are composite nanostructures consisting of nanogold shells directly coated on the Fe3O4 core surface. Therefore, it has a structure compared to most known multimode imaging probes. Stable, adjustable particle size, etc. Most importantly, it combines all the characteristics of nano-gold spheres and nano-ferric oxides. It can be used as a probe for dual-mode imaging of nuclear magnetic imaging and photoacoustic imaging, so it is a combination. A dual-mode imaging probe with multiple physical properties and complementary shortcomings. In addition, when we modified the Integrin v3 monoclonal antibody and FITC fluorescent dye on the surface of Fe3O4@Au, it can also selectively enter U-87 MG cells with high expression of Integrin v3, and fluorescence imaging of U-87 MG cells.

In summary, the bio-modified Fe3O4@Au nanoparticles synthesized by us can be used as a multifunctional probe for imaging tumor cells by nuclear magnetic and photoacoustic targeting specific cells. Due to its wide application in tumor imaging. Potential, this new imaging probe will be of great significance in promoting the development of clinical diagnosis of tumors.

## Materials and method

FeCl2·4H2O, FeCl3·6H2O, ammonia water (25%~28%), NaCl, KCl, sodium citrate tetrahydrate was purchased from Guangdong Chemical Reagent Company; HAuCl4·4H2O was purchased from Sinopharm Chemical Reagent Co., Ltd.; SHC11H22CO2H (11-MUA, 95%) was purchased from BEHRINGER Reagent Co., Ltd.; mPEG-SH (MW5000) was purchased from Jenkem Technology Co., Ltd.; Integrin v3 monoclonal antibody was purchased from Santa Cruz Biotechnology Company of the United States; FITC, NHS, EDC was purchased from Sigma- USA Aldrich Company; The above reagents are of analytical grade unless otherwise specified. The whole experiment uses deionized water with an impedance of >18 M cm.

Place FeCl3·6H2O in a four-reaction flask system, measure 100 mL of three distilled water, add FeCl3·6H2O to the reaction flask, protect with nitrogen, and add FeCl2·4H2O while stirring vigorously. The inlet is added dropwise to the reaction system. When the pH of the solution rises to about 9.0, stop adding ammonia. Continue the reaction for 15 min. Heat to 80 ° C and mature for 30 min.

Preparation of nano-Fe3O4@Au. Take 20 mL of Fe3O4 solution for magnetic separation and wash it with sodium citrate solution (0.2 g sodium citrate in 100 mL of trihydrated water) for 3 times, and finally dissolve in 100 mL of the same concentration of citric acid. In the sodium solution, placed in an ultrasonic machine for several hours of ultrasound. Place the mixture after sonication in a flask and slowly stir while heating to 70 ° C. Slowly add HAuCl 4 solution while stirring vigorously, and stop heating after 1 h of reaction. Stir for 40 min. Stir the reaction.

Coupling of Integrin v3 with FITC. Add 0.1 mg/mL FITC in DMSO to 400L Integrin v3 monoclonal antibody in PBS. Mix well and protect from light at 4 °C in the refrigerator. After several hours, the reaction-forming Integrin v3 mAb-FITC was dialyzed several times with a 1 kDa semipermeable membrane (Millipore) until all unreacted free FITC was removed.

Functional modification of nano-Fe3O4@Au. Add 100L of 1 mg/mL mPEG-SH to 1 mL of nano-Fe3O4@ Au solution. After ultrasonic reaction for 30 min, add 100L 11-MUA solution to continue the number of ultrasonic reactions. Hour. Magnetic separation wash to remove unattached reagent molecules, then add EDC and NHS, and FITC-incorporated Integrin v3 monoclonal antibody. After 4 °C overnight in the dark, the magnetic separation is washed several times. Remove excess reagents that are not connected.

U87-MG human glioma cells and MCF-7 human breast cancer cells were cultured in EMEM medium and DMEM medium respectively. 10% fetal bovine serum and 1% penicillin streptomycin were added to the culture medium. Conditions are set to 37 ° C, 5% CO2, 95% air.

Incubate 500L cells with bio-modified Fe3O4@Au nano-solution in an incubator for 1 h, then gently rinse the cells with fresh medium to elute non-phagocytic nanoparticles. Blocked U87-MG cells were pre-treated The integrin v3 monoclonal antibody was blocked. The treated cells were imaged using a confocal microscope (LSM510/ConfoCor2) combined system (Zeiss, Germany). The excitation source was a 488 nm Ar ion laser.

## Results and discussion

In order to verify the successful synthesis of Fe3O4@Au nanoparticles, we characterized Fe3O4@Au nanoparticles. From the TEM photos of Fe3O4 and Fe3O4@ Au (Fig. 1(a) and (b)), Fe3O4@Au nanometers can be seen. The radius of the particles is larger than that of Fe3O4, and the dispersion of Fe3O4@Au nanoparticles is better than that of Fe3O4. The main reason is that Au is coated on the surface of Fe3O4 nanoparticles to form Fe3O4@Au nanoparticles, and the gold shell increases the stability of Fe3O4 nanoparticles. Fig. 2(a) is the absorption spectrum of Fe3O4 nanoparticles, gold nanospheres and Fe3O4@Au nanoparticles. The absorption peak of Fe3O4@Au nanoparticles is 526 nm, and the spectral characteristics are the same as those of nanogold spheres. Fig. Fe3O4@Au The absorption peak of the nanoparticles is slightly red-shifted (compared to the nano-gold sphere), probably due to the increase of the radius. In the surface Raman enhancement technique, Raman scattering can be enhanced when gold is adsorbed on the surface of the detector, and then Increase the sensitivity of Raman detection. As shown in Figure 2(b), the Raman signal of 4,4’-bipyridine is very weak, but its Raman is present when nano-gold spheres or Fe3O4@ Au nanoparticles are present on its surface. The signal is enhanced. However, Fe3O4 nanoparticles do not have this effect. Therefore, the Raman enhancement effect of Fe3O4@Au nanoparticles is derived from the gold shell on the surface. The above experiments show that Au has been coated on the surface of Fe3O4 core to form Fe3O4@Au nanometer. particle.

**Figure 1.**
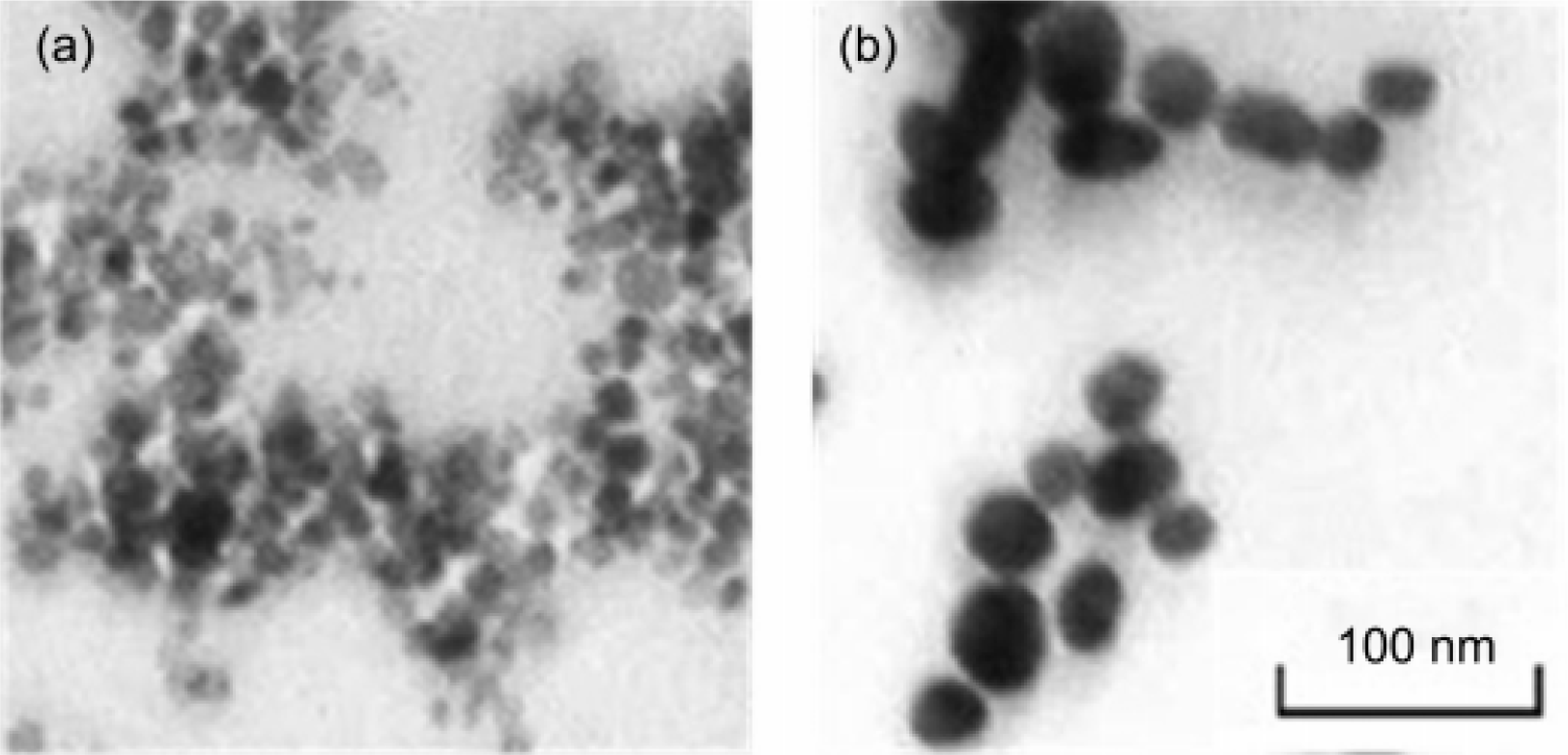
TEM analysis of Fe nanoparticles

Nuclear magnetic imaging is a non-destructive clinical imaging technique. In order to prove that Fe3O4@Au nanoparticles can be used as probes for nuclear magnetic imaging, the image enhancement effects of Fe3O4@Au and Fe3O4 nanoparticles were detected by nuclear magnetic T2 sequence. Figure 3(a) shows with the increase of [Fe] concentration (0, 0.1, 0.2, 0.4, 0.8, 1 mmol/L), the images of Fe3O4@Au and Fe3O4 nanoparticles become darker. The enhancement effect of Fe3O4@Au nanoparticles is slightly better than that of Fe3O4 nanoparticles. Poor. The transverse relaxation time of Fe3O4@Au nanoparticles is 26.53 m(mol/L) 1 s1, and the transverse relaxation time of Fe3O4 nanoparticles is 36.67 m(mol/L) 1 s1. Other Fe3O4/Au hybrid nanoparticles reported in the literature (such as dumbbell-shaped Au-Fe3O4 [13] and iron oxide/Au hybrid nanoparticles [7] connected by organic matter), Fe3O4@ at the same [Fe] concentration Au has similar nuclear magnetic contrast enhancement effects. In addition, due to the Au shell, the relaxation time of Fe3O4@Au nanoparticles is smaller than that of Fe3O4 nanoparticles. However, some commercial T2 weight nuclear magnetic contrast agents (such as dextran package) Compared to nano-iron oxide), Fe3O4@Au nanoparticles have Good stability properties and can be modified, the stability of these properties mainly from gold shell and easily modifying.

**Figure 2.**
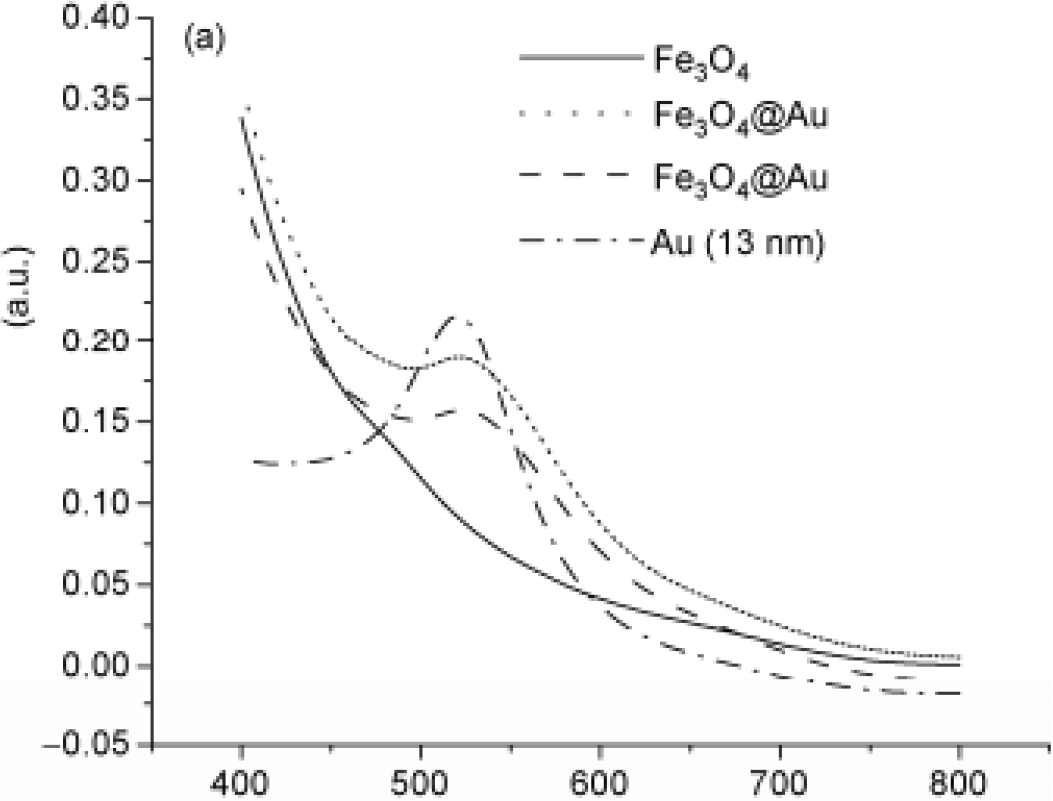
Releasing kinetic analysis

Photoacoustic imaging is a new type of non-destructive medical imaging technology that has just been developed in recent years. This imaging mode combines the high selectivity of optical imaging with the high resolution of ultrasound imaging [14~18]. Gold nanoparticles have been proven very It is suitable as a probe for photoacoustic imaging. In order to prove that Fe3O4@Au nanoparticles can be used as probes for photoacoustic imaging, the photoacoustic signals of Fe3O4@Au nanoparticles at different concentrations are measured. As shown in Fig. 3(b), With the increase of [Au] concentration (0.1, 0.2, 0.4, 0.6, 0.8, 1 mmol/L), the photoacoustic signal of Fe3O4@Au nanoparticles is continuously enhanced, and it turns out that Fe3O4@Au nanoparticles can be used as photoacoustic imaging. Probe. The above experiments show that Fe3O4@Au nanoparticles can enhance both nuclear magnetic and photoacoustic signals, so it can be used as a dual-mode probe for nuclear magnetic and photoacoustic imaging.

**Figure 3.**
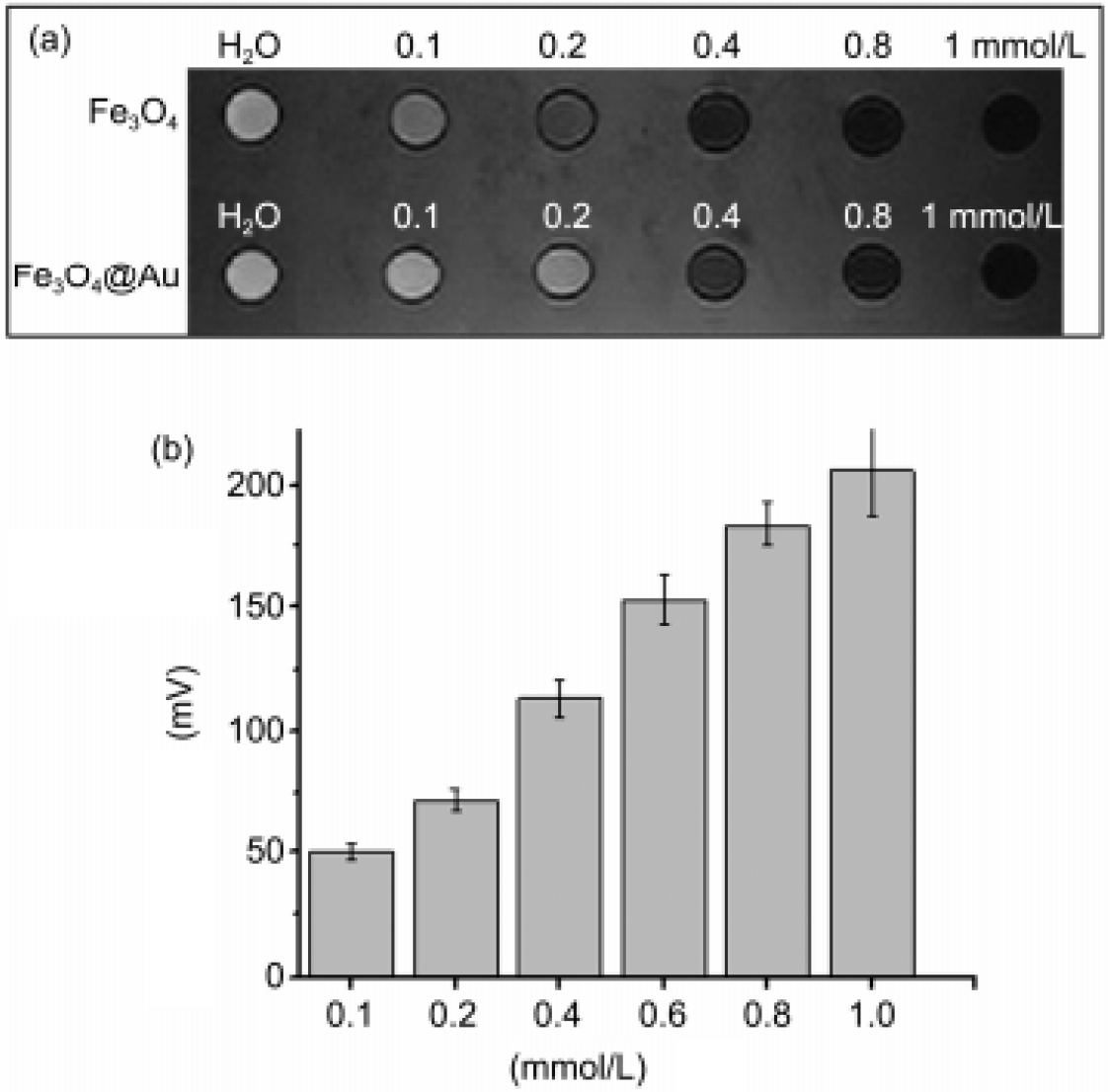
Cytotoxicity analysis of Fe nanoparticles

In order to make the Fe3O4@Au nanoparticles have tumor cell targeting function, we modified the Integrin v3 monoclonal antibody on the surface. At the same time, the Integrin v3 monoclonal antibody-conjugated Fe3O4@Au nanoparticle was labeled with FITC. The particles were observed by confocal microscopy to see if the nanoparticles entered the tumor cells. U87-MG tumor cells overexpressed v3 on the surface of the cell membrane. However, there were only a few v3 on the surface of MCF-7 tumor cells. Bio-modified Fe3O4@ Au nanoparticles were co-incubated with U87-MG and MCF-7, respectively. As shown in Figure 4, in the confocal microscopy images, there was a strong fluorescent signal in the cultured U87-MG tumor cells, whereas MCF-7 tumor cells No fluorescence signal was found in the U87-MG tumor cells of the control group. The confocal experiments of the cells demonstrated that the bio-modified Fe3O4@Au nanoparticles have selective targeting to U87-MG tumor cells, and the bio-modified Fe3O4@Au nanometer. Particles may be used in tumor diagnosis in the future.

**Figure 4.**
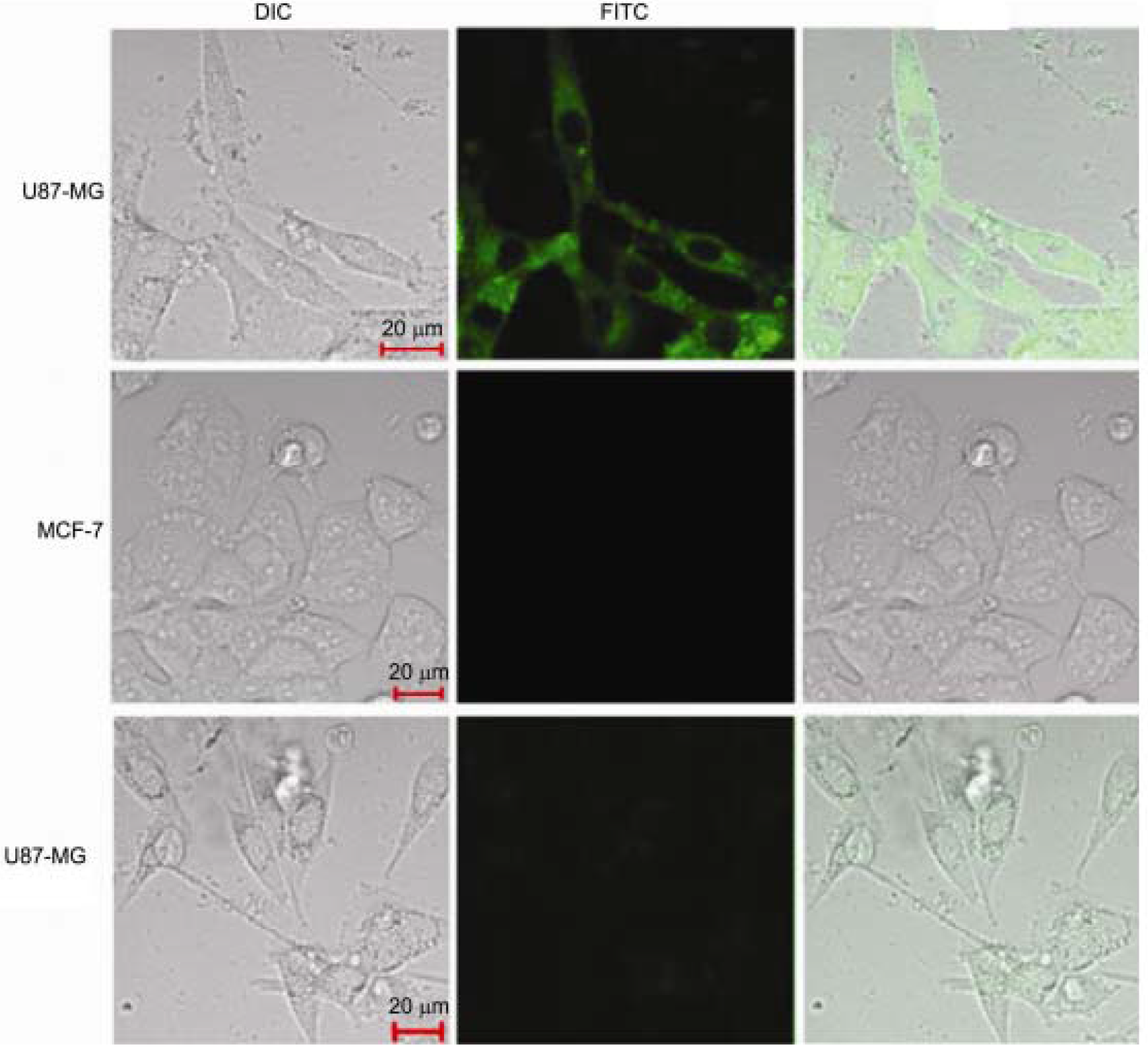
Cell uptake analysis by fluorescent microscopy.

Confocal fluorescence experiments demonstrated that Fe3O4@Au nanoparticles can be effectively targeted to U87-MG tumor cells. Different numbers of U87-MG tumor cells were labeled with Fe3O4@Au nanoparticles, using nuclear magnetic imaging system and photoacoustic microscopy, respectively. The imaging system images four different numbers of cell samples (approximately 1.2 × 105, 3.6 × 106, 7.4 × 106, 2.5 × 107) (Figure 5). As the number of labeled cells increases, the nuclear magnetic signal and light of the T2 sequence The acoustic signal is enhanced, and the minimum detection amount of the two imaging pairs is about 3×106. The above experiments fully prove that when the U87-MG cells are labeled by Fe3O4@Au nanoparticles, they can be simultaneously tracked by nuclear magnetic/photoacoustic imaging. In in vivo tumor diagnosis, nuclear magnetic imaging can be used as a systemic scan to determine the location of the tumor; photoacoustic imaging can analyze the physiological parameters of the tumor microenvironment, such as blood oxygen saturation, pH. Because photoacoustic imaging has submicron level Resolution, which can obtain morphological information of tiny areas of the tumor (such as micro-new blood vessel structure), and then determine the nature of the tumor from a morphological point of view. The decorated Fe3O4@Au nanoparticles can be actively targeted to U87-MG tumor cells, and the labeled U87-MG tumor cells can be tracked by nuclear magnetic/photoacoustic imaging. The combined imaging of nuclear magnetic and photoacoustic can be multi-parameter, multi-scale Multi-dimensional reflection of tumor information. Bio-modified Fe3O4@Au nanoparticles will greatly promote the application of nuclear magnetic and photoacoustic dual-mode imaging in clinical practice.

## Conclusion

In this paper, bio-modified Fe3O4@Au nanoparticles were successfully synthesized. The bio-modified Fe3O4@Au nanoparticles can simultaneously enhance the nuclear magnetic signal and photoacoustic signal of T2 sequence. This probe has selective targeting to U87-MG tumor cells. U87-MG tumor cells after novel probe labeling can be simultaneously tracked by nuclear magnetic and photoacoustic imaging. Bio-modified Fe3O4@Au nanoparticles will be the bridge between nuclear magnetic and photoacoustic dual-mode imaging. Nuclear magnetic/photoacoustic dual-mode imaging will be able to obtain more Multiple tumor morphology and physiological information, nuclear magnetic/photoacoustic dual-mode imaging based on Fe3O4@Au nanoparticles will play an important role in the clinical diagnosis of tumors.

